# Differential oxidative and pro-apoptotic response of cancer and normal cells to an anti-inflammatory agent CLEFMA

**DOI:** 10.1101/2021.06.02.446782

**Authors:** Kaustuv Sahoo, Vibhudutta Awasthi

## Abstract

Selective killing of cancer cells by chemotherapy has been an age old challenge, but certain unique features of cancer cells allow discriminatory response between cancer and normal cells. The objectives of this study was to investigate pro-oxidant and apoptotic effects of CLEFMA, an anti-inflammatory compound with anticancer activity, in lung cancer cells versus normal lung fibroblasts and to establish its maximum tolerated dose (MTD) in mice. We found that CLEFMA preferentially induced reactive oxygen species (ROS)-mediated apoptosis in H441, H1650 and H226 cancer cells, but spared normal CCL151 and MRC9 fibroblasts. Immunoblotting studies revealed that CLEFMA-induced apoptosis is associated with p53 phosphorylation in cancer cells which was not observed in CLEFMA treated normal fibroblasts. CLEFMA showed no effect on NF-κB p-65 expression in the normal lung fibroblasts, whereas its translocation to nucleus was inhibited in cancer cells. Furthermore, CLEFMA treatment also inhibited the DNA-binding activity of NF-κB p65 in H441cancer cells, but not in normal CCL151 cells. Preclinical toxicology studies in CD31 mice showed that CLEFMA was not toxic when injected daily for 7 days or injected weekly for 4 weeks. Based on survival data, MTD of CLEFMA was estimated as 30 mg/kg bodyweight. We conclude that CLEFMA exploits the biochemical differences in cancer and normal cells and selectively induces ROS in cancer cells. Secondly, CLEFMA can be safely administered in vivo because its known dose necessary for *in vivo* efficacy as anti-inflammatory and anti-tumor agent (0.4 mg/kg) is 75 times lower than its MTD.

## INTRODUCTION

Chemotherapy is widely used for treatment of cancer, either alone or in combination with other radical therapies such as surgery and radiation. For instance, over 85% of patients diagnosed with non-small cell lung carcinoma (NSCLC) depend on systemic chemotherapy as part of the overall management [1]. Chemotherapy has definite palliative benefits in symptomatic relief and improvement in quality of life, even if it fails to prolong survival in patient population in meaningful fashion. Since anti-cancer therapies are cytotoxic by their very design, a major goal in oncology is to develop drug therapies exhibiting selectivity for cancer cells with minimum toxicity to normal cells. Chemotherapeutic choices in lung cancer depend on histological typing, but highly toxic platinum-based regimens continue to remain important. Novel molecularly-targeted drugs (e.g., erlotinib) show promise, but are riddled with problems of intra-tumoral/inter-patient molecular homogeneity, development of resistance, etc. [2-6]. Even if tumor heterogeneity and emergence of resistance are overcome, targeted therapies succeed only when patient population is segregated into target-selective subpopulations in a timely fashion [7]. Not surprisingly, a consensus is building that targeted therapies do not dramatically change clinical outcome for most patients [8, 9].

Cancer cells find it easy to develop resistance against drugs that are dependent only on influencing single molecule or pathway. In most lung cancer cells, the molecular pathways of cell death are mutated, resulting in a resistance against drugs. Small molecules that can affect multiple critical components, and that may induce alternative cell death mechanisms, may provide better outcomes than the agents that target single step in signaling pathways. The selectivity of such therapies is dependent on affecting biochemical differences between cancer cells and normal cells. An example to this effect was an early observation that cancer cells exhibit enhanced aerobic glycolysis [10]. Because of the resultant high metabolic turnover cancer cells are more oxidatively stressed and generate larger amounts of reactive oxygen species (ROS) than the normal cells. In order to survive this oxidative stress and keep ROS at benign levels, cancer cells have upregulated antioxidant system that includes superoxide dismutase (SOD), glutathione peroxidase (GPX), catalase, etc. However, if ROS exceeds the tolerable level, cancer cells undergo programmed cell death. In contrast, normal cells demonstrate higher threshold to oxidative stress [11, 12]. The exploitation of ROS as a trigger for cancer cell death is not entirely novel because radiotherapy survives on this principle. However, identification of pharmacologic choices for selective induction of ROS in cancer cells is novel [13, 14].

CLEFMA or 4-(3, 5-bis (2-chlorobenzylidene)-4-oxo-piperidine-1-yl)-4-oxo-2-butenoic acid is a chalcone derivative that exploits the unique susceptibility of cancer cells towards oxidative stress [15, 16]. This characteristic of CLEFMA is believed to originate from its unique mechanism of the 26S proteasome inhibition [17]. Its ability to induce apoptosis in lung adenocarcinoma model has been reported [18] and its anti-proliferative efficacy in NSCLC cell lines is comparable to some of the well-established anticancer drugs [16]. More recently, Yang et al. reported that jun-N-kinase and p38 signaling is important for CLEFMA-induced apoptosis osteosarcoma cells [19]. Here, we provide evidence that the selectivity of CLEFMA against lung cancer cells in comparison to that in normal lung fibroblasts is dependent on discriminatory ROS generation in cancer cells. We also performed a preclinical toxicological study to estimate maximum tolerated dose (MTD) of CLEFMA.

## MATERIALS AND METHODS

### Materials

CLEFMA was synthesized in-house and analyzed for purity by the methods detailed elsewhere [15]. Celecoxib was obtained as a gift sample from Dr. CV Rao (Biomedical Research Center, OUHSC, OK, USA). The mouse monoclonal antibodies against survivin, B-cell lymphoma 2 (Bcl2), B-cell lymphoma-extra large (Bcl-xL) and cellular inhibitor of apoptosis protein-1 (cIAP1) were supplied by Santa Cruz Biotechnology (Santa Cruz, CA, USA). Rabbit polyclonal antibodies against cleaved poly (ADP-ribose) polymerase or PARP, total PARP, caspase-3, cleaved caspase-3, phospho-p65-NF-κB, p65, and phospho-p53 were purchased from Cell Signaling Technology (Boston, MA, USA). Anti-β-actin antibody was obtained from Sigma-Aldrich (St. Louis, MO, USA). Secondary antibodies against IgG of various species were from Santa Cruz Biotechnology and Cell Signaling Technology. We obtained catalase (CAT) from Worthington Biochemical Corp. (Lakewood, NJ, USA), superoxide dismutase (SOD) from Sigma-Aldrich Earlier, and NAC from Enzo Life Sciences (Plymouth Meeting, PA, USA).

### Cell lines and cell culture

Human NSCLC cell lines NCI-H441, H226, and H1650 were obtained from American Type Culture Collection (ATCC, Manassas, VA, USA). The cells were maintained at 37°C with 5% CO_2_ in RPMI 1640 medium (Invitrogen, Carlsbad, CA, USA) supplemented with 10% heat-inactivated fetal bovine serum (FBS) and gentamicin at 50 μg/mL concentration. Similarly, normal lung fibroblasts (CCL151 and MRC9 from ATCC) were grown in EMEM medium supplemented with 15% heat-inactivated fetal bovine serum.

### Cell viability assay

Effect of CLEFMA on cell viability of cancer and normal cells was determined by 3-(4,5-dimethylthsiazol-2-yl)-2,5-diphenyltetrazolium bromide (MTT) assay, as previously described [18]. In brief, approximately 5,000 cells per well were allowed to grow for 24 h in a 96-well plate at 37 °C, followed by treatment with CLEFMA. At the end of 48 h incubation, 20 μL of MTT solution (5 mg/mL) was added into each well. After 2 h of incubation, the content in each well was dissolved in lysis buffer containing 20% SDS and 50% dimethylformamide and the color was quantified at 570 nm using a microplate reader (BioTek, Winooski, VT, USA).

To compare the toxicity of CLEFMA and celecoxib on normal cells, we tested the viability of normal lung fibroblast CCL151 cells that were treated at twice the IC_50_ levels of CLEFMA and celecoxib for H441 cells, respectively. IC_50_ values of CLEFMA and celecoxib for H441 cells was established as 6 µM and 25 µM prior to these experiments.

### Cell Proliferation

Cell proliferation was examined by 5-bromo-2-deoxyuridine (BrdU) cell proliferation enzyme-linked immunosorbent assay (ELISA) according to the kit manufacturer’s protocol (Life Technologies, Grand Island, NY, USA). The cells were incubated in presence of various concentrations of CLEFMA (0 – 10 µM). After 48 h incubation at 37 °C, the cells were labeled with 100 µl of 1:100 diluted BrdU for 12 h at 37 °C. After removing the labeling solution, the cells were gently washed two times with PBS fixed with 100 µl of 70% alcohol. The fixing was allowed to occur for 30 min at room temperature and the fixed cells were washed three times with PBS. Each well was incubated with 100 µl of the anti-BrdU-peroxidase antibody solution (1:1,000) for 90 min at room temperature. After several washing steps, 100 µl tetramethylbenzidine substrate solution was added per well for 20 min and the reaction was stopped with 50 µl of 1M H_2_SO_4_. The incorporation of BrdU into DNA was quantified by measuring absorbance of the chemiluminescent substrate using a BioTek 96-well multiscanner.

### Reactive oxygen species (ROS) measurement

Induction of intracellular ROS in H441 and CCL151 cells upon CLEFMA treatment in presence and absence of ROS scavengers was monitored by a fluorescence-based OxiSelect assay (Cell Biolabs, Inc., San Diego, CA, USA) as described previously [16]. Briefly, a cell-permeable fluorogenic probe 2′, 7′-dichlorodihydrofluorescin diacetate (DCFH-DA) was added to the cells cultured in 96-well plates and the plate was incubated for 30–60 min. The wells were washed twice with PBS and treated with CLEFMA in presence or absence of NAC (1 mM), CAT, (1000 U), and SOD (500 U). Earlier, non-toxic concentrations of CAT, SOD, and NAC were established in H441 cells. After the incubation period, the medium was removed and the cells were gently washed 2– 3 times with PBS. The fluorescence was measured at λ_Ex_ 480 nm/ λ_Em_ 530 nm using Cytoflour 2300 (Millipore, Billerica, MA, USA).

### Fluorescence imaging of ROS

For visualization of intracellular ROS, cells were stained with Image-iT LIVE Green ROS detection kit (Life Technologies) according to the protocol provided by the manufacturer. All dilutions and washings were done with Hank’s balanced salt solution with calcium and magnesium (HBSS/Ca/Mg). Briefly, the cells were treated with CLEFMA for 24 h. After the incubation period, the medium was removed and the cells were gently washed 2–3 times with HBSS followed by incubation with carboxy-H_2_DCFDA in PBS for 30 min at 37 °C. This was followed by counterstaining the cells with Hoechst 33342 which was added at a final concentration of 1 μM to the carboxy-H_2_DCFDA staining solution during the last 5 min of the incubation period. The cells were then imaged under fluorescent microscope equipped with Olympus DP70 camera (Melville, NY, USA) at λ_Ex_/λ_Ex_ of 350/461 nm for Hoechst 33342 and 495/529 nm for carboxy-H_2_DCFDA.

### NF-κB DNA-binding

The effect of CLEFMA on NF-κB p65-DNA interaction was assessed in the nuclear fraction of H441 and CCL151 cells by TransAM NF-κB p65 kit according to the manufacturer’s instructions (Active Motif, Carlsbad, CA, USA). A specific NF-κB inhibitor, BAY 11-7082 (Cayman Chemical, Ann Arbor, MI, USA), was used at 10 μM as a positive control. Briefly, cells were transfected with the NF-κB firefly luciferase reporter plasmid pGL4.32 [luc2P/NF-κB-RE/Hygro, Promega, Madison, WI, USA; kindly provided by Dr. Kelly Standifer, University of Oklahoma Health Sciences Center (OUHSC)]. For ensuring successful transfection, the control wells of H441 cells were transfected with a plasmid DNA construct expressing enhanced green fluorescent protein (pHYG-EGFP; Clontech, Mountain View, CA, USA) and observed under Leica DM4000B fluorescent microscope.

### Western blotting

CLEFMA-treated cancer and normal cells were incubated on ice for 30 min in 0.5 mL of ice-cold lysate buffer consisting of 10% NP-40, 5 M NaCl, 1 M HEPES, 0.1 M ethyleneglycoltetraacetic acid, 0.5 M ethylenediaminetetraacetic acid, 0.1 M phenylmethylsulfonyl fluoride, 0.2 M sodium orthovanadate, 1 M NaF, 2 μg·mL^− 1^ aprotinin and 2 μg·mL^− 1^ leupeptin. The protein was extracted by homogenization using a probe sonicator and centrifugation at 16,000× g at 4°C for 10 min. The proteins were fractionated by SDS-PAGE, electrotransferred on to nitrocellulose membranes, and blotted with respective primary antibodies, followed by an HRP-conjugated secondary antibody. The final detection was performed by enhanced chemiluminescence with SuperSignal West Femto reagent (Thermo Scientific, Rockford, IL, USA). Equal protein loading was confirmed by stripping the blot and re-probing it with 1:1,000 anti-actin antibody.

### Maximum tolerated dose (MTD)

All animal care and experimental protocols were reviewed and approved by the Institutional Animal Care and Use Committee at the OUHSC. For acute toxicological study, CD31 mice (6 weeks old) were obtained from Harlan Laboratories (Indianapolis, IN, USA). Before initiating the experiment, we allowed the mice to acclimatize for at least 5 days. The mice were allowed *ad libitum* food and water during the study. CLEFMA solution was prepared in PEG_400_-saline and sterilized by filtration through 0.22 µm filter. It was administered by intraperitoneal injection once daily (1-week/daily-dosing study) in 100 µl volume at doses of 0, 1, 10, 50, 100, 200, and 400 mg/kg bodyweight. Each dose was tested in four mice (2 male + 2 female). The mice were monitored daily for weight and clinical signs up to 7 days of injection. All mice (including those which died during the test) were subjected to necropsy. Whole blood, serum, and urine were submitted to IDEXX Preclinical Research (Sacramento, CA, USA) for analyses. The MTD of CLEFMA was taken as the dose above which there was 10% body weight reduction. MTD was also calculated based on the dose above which there was one or more incident of death. Further, a safe maximum dose of CLEFMA based on its MTD was administered by intraperitoneal injection once every week for four weeks (4-week/weekly-dosing study) to assess its 4-week toxicity.

### Statistical analysis

All values in figures and text are presented as mean ± SEM. Differences among group means were assessed by ANOVA (analysis of variance). Survival data were analyzed by Kaplan-Meier Mantel-Cox Logrank test. Differences were considered statistically significant for p-value of < 0.05. GraphPad Prism software 6.0 (GraphPad Software, Inc. La Jolla, CA, USA) was used to perform all statistical analyses.

## RESULTS

### CLEFMA induces cell death selectively in cancer cells

We studied the effect of CLEFMA on viability and DNA turnover of three NSCLC cell lines and one normal lung fibroblast cell line. As shown in **Figure 2**, CLEFMA inhibited the viability and proliferation of H441, H1650, and H226 cancer cells in a dose-dependent manner. The IC_50_ values for CLEFMA were calculated on the basis of this viability data in Figure 2A. The IC_50_ were 6.1 μM in H441 cells, 5.86 μM in H1650 cells and 7.1 μM in H226 cells. We found that normal lung CCL151 fibroblasts were not sensitive to CLEFMA-induced cell death at the concentrations tested. We also compared the effects of CLEFMA and celecoxib on the viability of normal lung CCL151 fibroblasts. Treatment of normal lung fibroblasts CCL151 with CLEFMA and celecoxib showed significant differences in the effect of two drugs (**Figure 3**). Whereas celecoxib significantly affected the viability of normal lung fibroblasts at dose level of 50 µM, CLEFMA was not found to have any significant effect on the viability of normal cells even at twice the dose of its IC_50_ value (12 µM).

**Figure 1:**
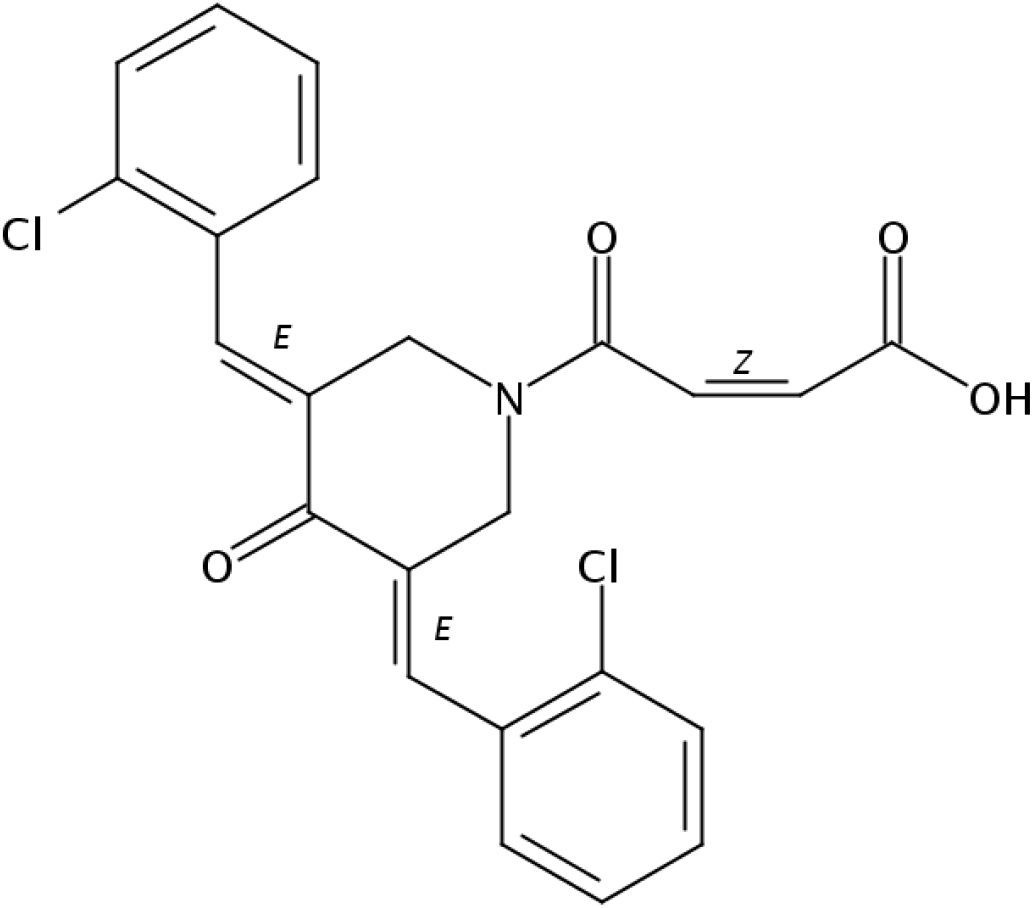
Chemical structure of CLEFMA.

**Figure 2:**
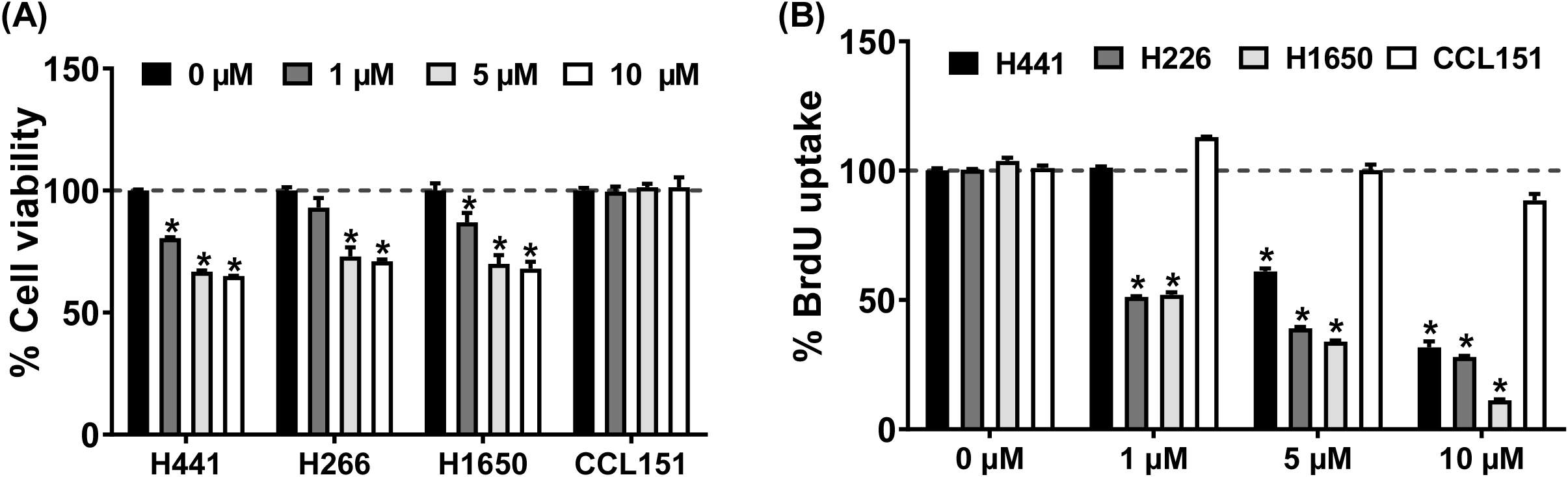
CLEFMA inhibits viability and proliferation of cancer cells (H441, H226, and H1650) and not of normal fibroblasts (CCL151).

**Figure 3:**
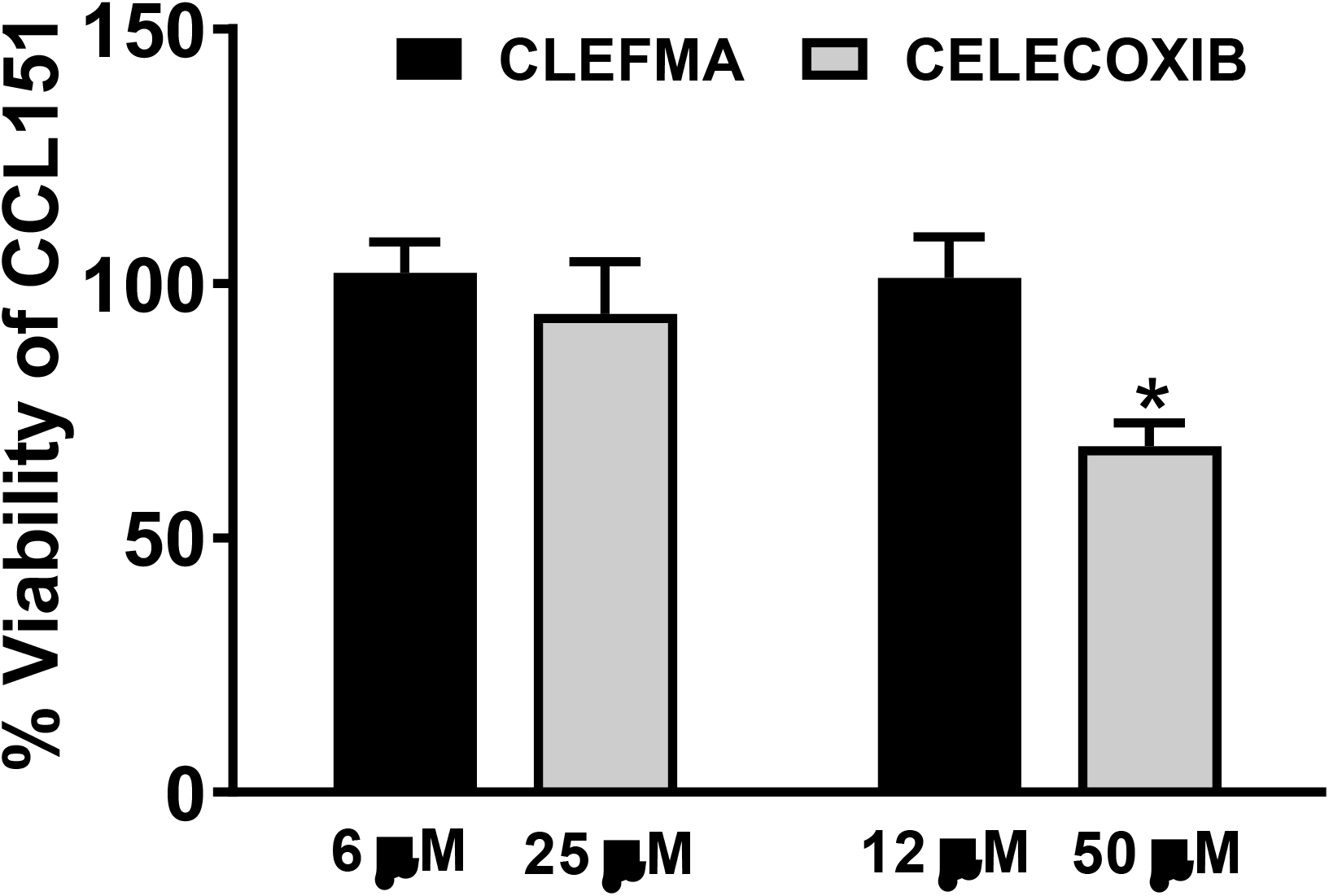
Comparative cytotoxicity of CLEFMA and celecoxib in normal lung fibroblasts CCL151. Cell viability was determined at twice the IC_50_ value of celecoxib and CLEFMA in H441 cells (50 µM and 12 µM, respectively; *p < 0.05 compared to control).

### CLEFMA induces ROS in cancer cells, but not in normal cells

As shown in **Figure 4A**, CLEFMA treatment generated ROS in lung adenocarcinoma H441 cells. CLEFMA-induced ROS generation was suppressed by a mixture of antioxidant principles CAT, SOD, and NAC. Importantly, CLEFMA did not increase ROS production in normal lung fibroblasts CCL151 (**Figure 4B**). These results were further confirmed by fluorescence imaging of the cells by labeling the cells with a fluorescent marker, H_2_DCFDA, signifying the presence of ROS (**Figure 5**). CLEFMA specifically produced ROS in all three cancer cells but spared the two normal lung fibroblast cell lines CCL151 and MRC9. Together, these results suggest that CLEFMA selectively induces ROS in cancer cells.

**Figure 4:**
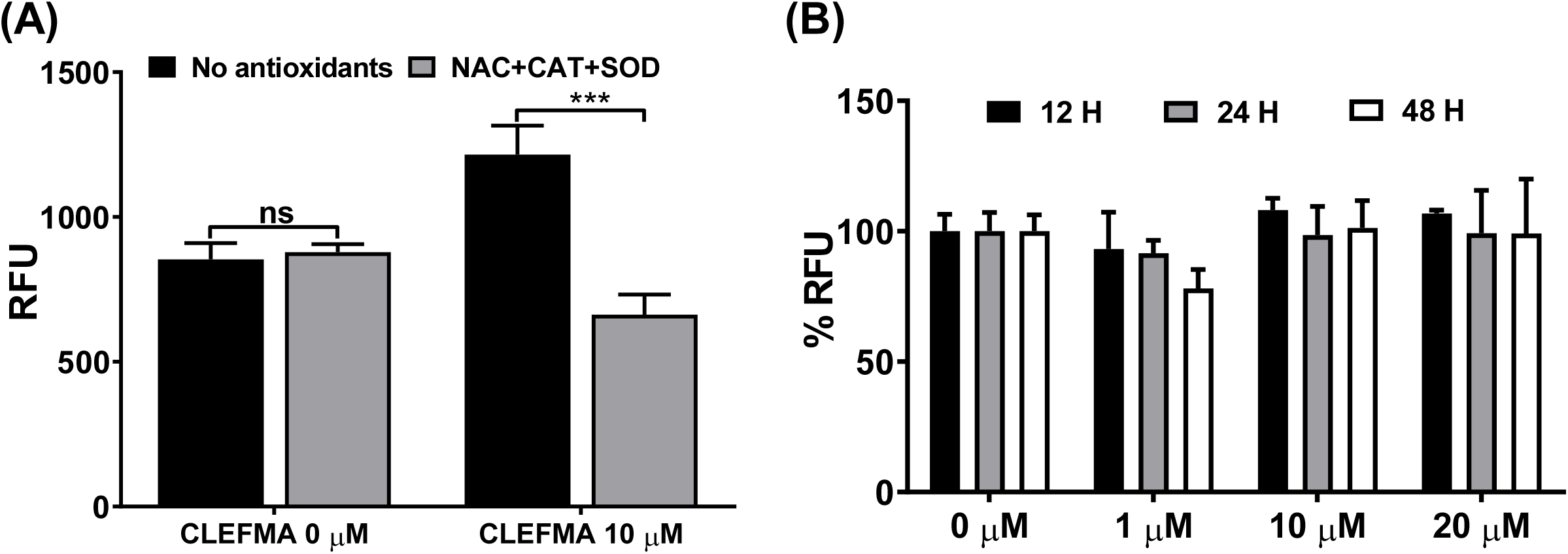
CLEFMA treatment generates ROS in cancer cells, but not in normal lung fibroblasts. (A) Effect of CLEFMA on induction of ROS in H441 lung cancer cells and normal lung fibroblasts CCL151 in presence or absence of antioxidants (*p <0.05 compared with control). (B) CLEFMA did not induce ROS in normal lung fibroblasts CCL151 at all concentrations of CLEFMA tested. ROS was measured in terms of relative fluorescence unit (% RFU) as compared to control cells which were not treated with CLEFMA.

**Figure 5:**
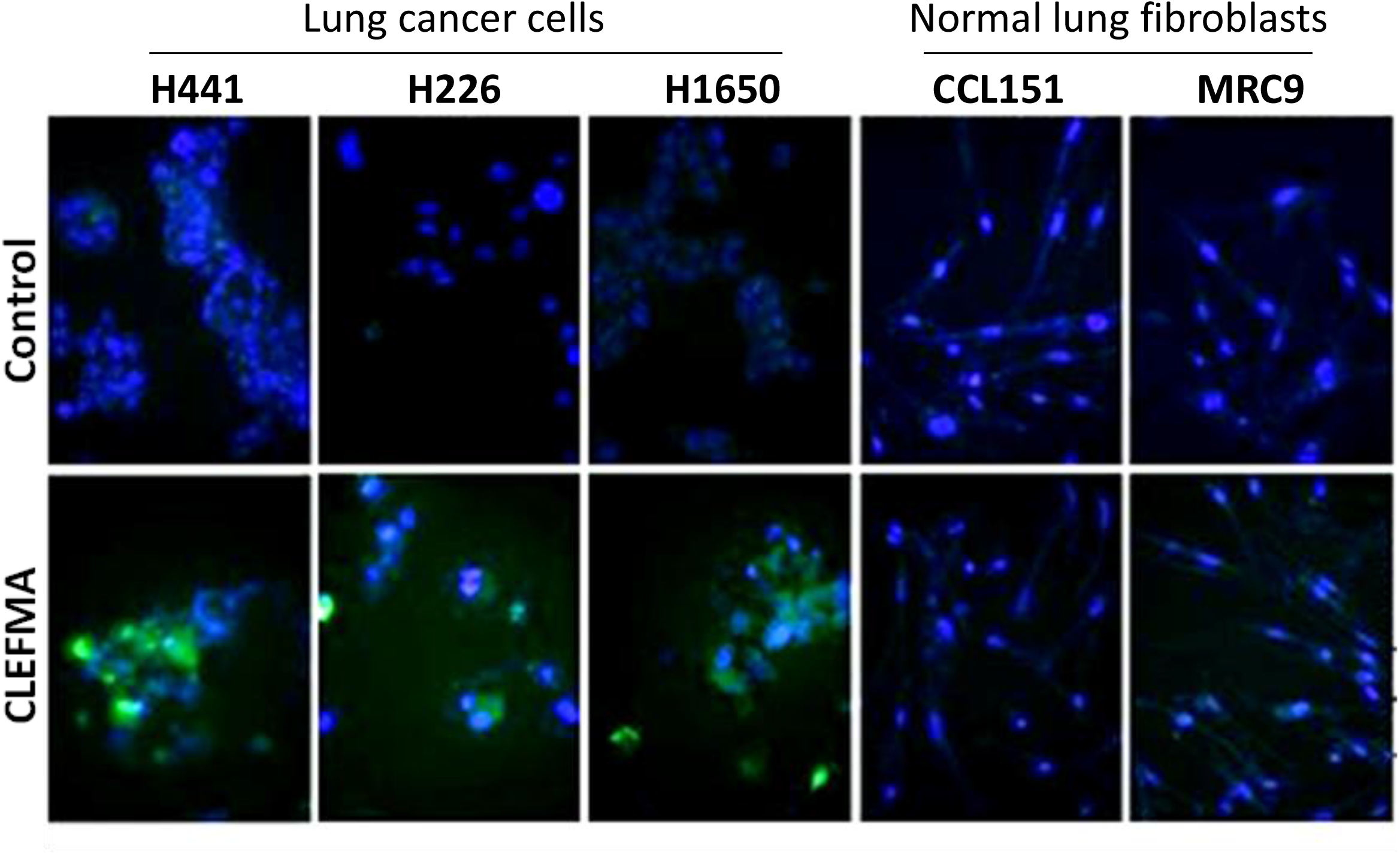
Fluorescence micrographs of cancer cells (H441, H226, and H1650) and normal fibroblasts (CCL151 and MRC9) showing differential ROS activity. The cells were stained with Image-iT LIVE Green Reactive Oxygen Species Detection kit. Blue represent nuclear staining and green punctuates track the ROS generated. Control cells received vehicle DMSO treatment whereas treated cells received CLEFMA at 10 μM concentration.

### CLEFMA induces apoptotic markers in cancer cells, but not in normal cells

To confirm that the CLEFMA-induced cell death in cancer cells is mediated by ROS, we assessed apoptotic markers in CLEFMA-treated cancer cells and normal cells. CLEFMA treatment significantly down-regulated the expression of anti-apoptotic molecules cIAP1, Bcl-xL, Bcl2, and survivin in cancer cells, without affecting their levels in normal cells (**Figure 6A**). Further, CLEFMA increased the accumulation of pro-apoptotic molecules: cleaved PARP and caspase-3 in cancer cells, but such changes were not observed in normal cells (**Figure 6A**). ROS-induced caspase-dependent apoptosis has been shown to be mediated by tumor suppressor p53 [20]. Upon investigation of p53 status we found that CLEFMA induced the phosphorylation of p53 in cancer cells only (**Figure 6B**). It is noteworthy that all three NSCLC cell lines showed similar sensitivity to CLEFMA treatment. These results show that cancer cells are sensitive to CLEFMA-induced apoptotic cell death, but normal lung fibroblasts show resistance to CLEFMA-induced apoptosis.

**Figure 6:**
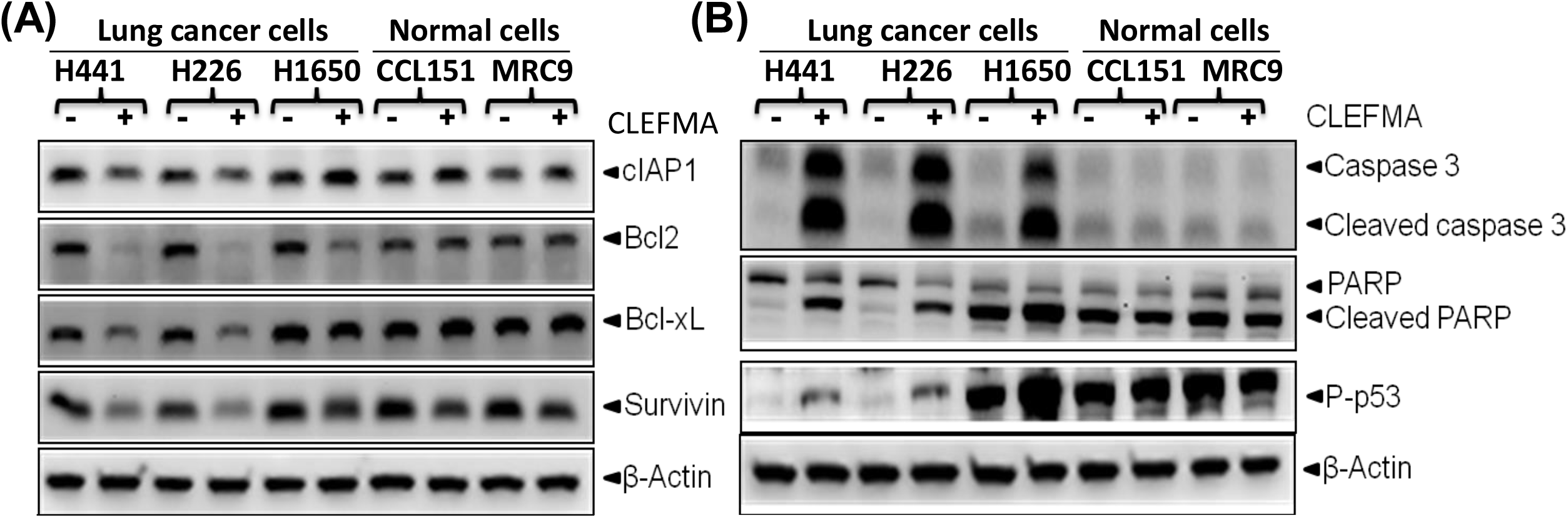
CLEFMA induces apoptosis selectively in cancer cells. (A) Effect of CLEFMA on anti-apoptotic proteins in cancer cells (H441, H226, and H1650) and normal fibroblasts (CCL151 and MRC9). (B) Effect of CLEFMA on phospho-p53 expression and cleavage of apoptotic markers caspase 3 and PARP in cancer cells (H441, H226, and H1650) and normal fibroblasts (CCL151 and MRC9). The protein extracts were analyzed by Western blotting.

### CLEFMA inhibits NF-κB (p65) in cancer cells, but not in normal cells

In our previous study, we have shown that CLEFMA inhibits NF-κB [18]. We assessed NF-κB status by immunoblotting for phosphorylated form of p65 subunit of NF-κB and by estimating its DNA-binding activity. Immunoblotting of the nuclear extracts demonstrated that CLEFMA significantly reduced the nuclear levels of the phospho-p65 subunit in cancer cells whereas, no such effect of CLEFMA on nuclear phospho-p65 was observed in the normal fibroblasts (**Figure 7**). These observations are suggestive of an interesting possibility that inhibition of NF-κB by CLEFMA is specific to cancer cells.

**Figure 7:**
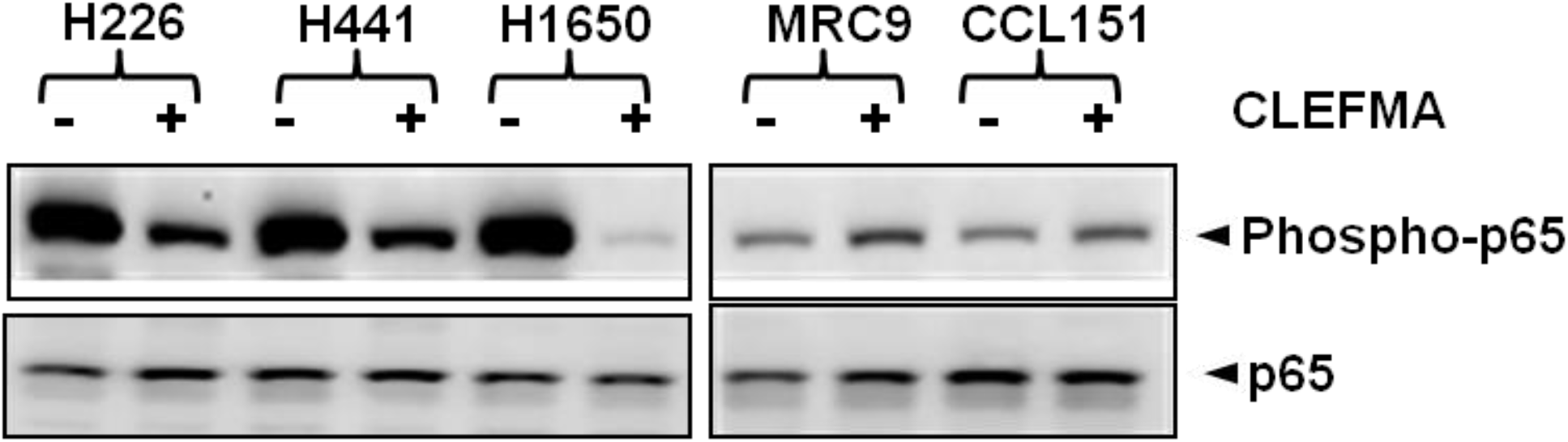
CLEFMA suppresses nuclear expression and activity of NF-κB in cancer cells, but not in normal fibroblasts. A representative blot showing that CLEFMA inhibited the nuclear translocation of NF-κB phospho-p65 in all cancer cells (H441, H1650, and H226), but had no such effect in normal fibroblasts (MRC9 and CCL151).

### Survival and MTD

**Figure 8A** shows the fraction of mice that survived after intraperitoneal administration of CLEFMA at various doses. There was a significant difference in survival curves between CLEFMA doses above 100 mg/kg and doses below 50 mg/kg. All the mice from dose groups that received 1 and 10 mg/kg of CLEFMA survived, but there was significant death associated with dose groups 100 mg/kg and above. Since only one incident of death was observed at 50 mg/kg dose and no death was observed at 10 mg/kg dose, the MTD_Survival_ for CLEFMA was estimated as average of the two i.e., 30 mg/kg bodyweight. The survival trend was found to be significant (p<0.05, Kaplan-Meier; Mantel-Cox Logrank test). Another criterion for MTD calculation was based on 10% or more reduction in bodyweight (MTD_10%BW_). There was significant reduction in body weight (more than 10% body weight reduction) in groups of mice which were administered with 50 mg/kg or more of CLEFMA (**Table 1**). Thus, on the basis of bodyweight reduction, the estimated MTD_10%BW_ was 50 mg/kg bodyweight.

**Figure 8:**
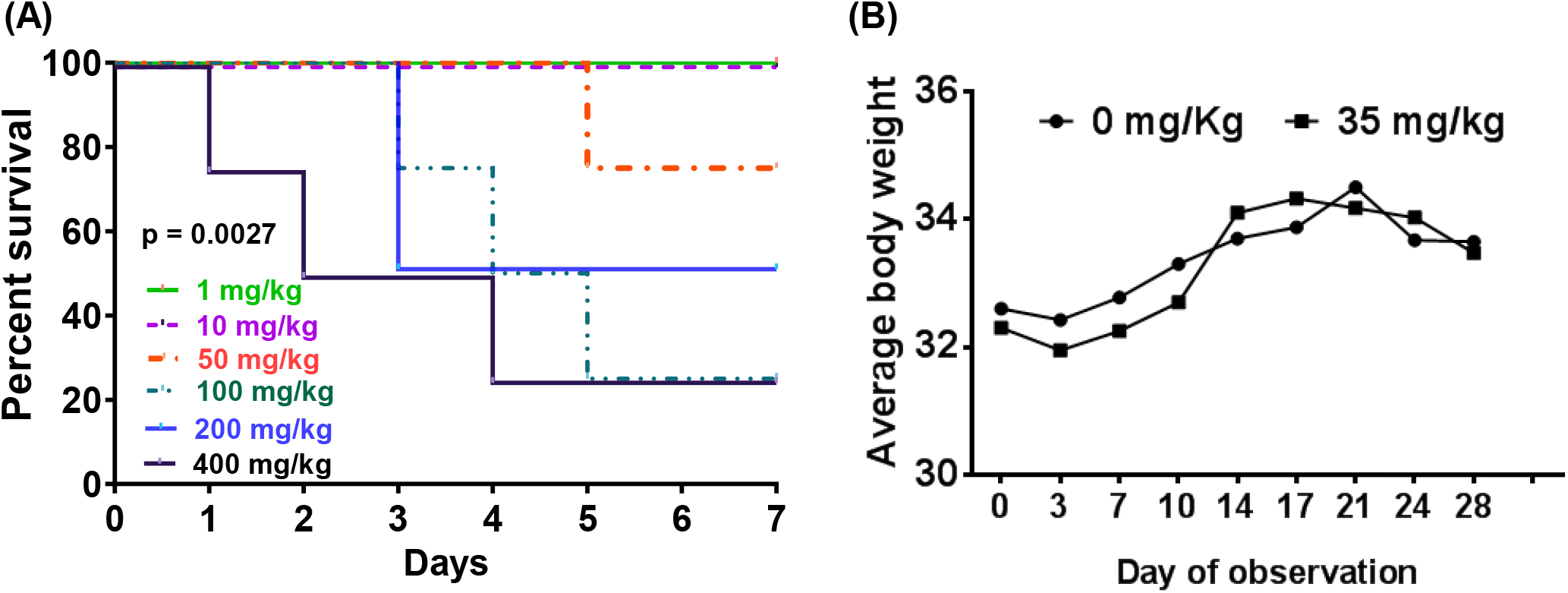
Effect of CLEFMA on the Kaplan Meier’s estimate of survival in CD31 mice. (A) Kaplan Meier survival curve of CLEFMA at different doses. Each group of CD31 mice had 2 animals/sex. The animals were administered six doses of CLEFMA by daily intraperitoneal injection for 7 days in a 1-week/daily-dosing study (p value of 0.0027 indicates significant trend for Logrank test). (D) Effect of CLEFMA on average body weight of mice in 4-week/weekly-dosing study. The animals were administered CLEFMA once every week for four weeks.

**Table 1.** Change in body weight of healthy CD31 mice that received CLEFMA.

| **CLEFMA dose**<br>**(mg/kg)** | **Number of mice showing more than 10% body weight reduction** |  |  |  |  |  |  |  |
| --- | --- | --- | --- | --- | --- | --- | --- | --- |
|  | **Day 0** | **Day 1** | **Day 2** | **Day 3** | **Day 4** | **Day 5** | **Day 6** | **Day 7** |
| **1** | 0 | 0 | 0 | 0 | 0 | 0 | 0 | 0 |
| **10** | 0 | 0 | 0 | 0 | 0 | 0 | 0 | 0 |
| **50** | 0 | 0 | 0 | 1 | 1 | 0 | 0 | 0 |
| **100** | 0 | 0 | 1 | 2 | 1 | 0 | 0 | 0 |
| **200** | 0 | 0 | 0 | 1 | 1 | 0 | 0 | 0 |
| **400** | 0 | 0 | 0 | 0 | 0 | 0 | 0 | 0 |

For four-week assessment of toxicity, we used a dose of 35 mg/kg which is close to the average of MTD_10%BW_ and MTD_Survival_. In this study, CLEFMA was administered on weekly basis. We observed that 35 mg/kg dose was safe because none of the mice showed any reduction in body weight (**Figure 8B**) and no incident of death was observed over the period of 4 weeks. Hematoxylin & Eosin stained sections of major organs (heart, liver, lung, and kidney) of CLEFMA treated and control mice are shown in **Figure 9**. We found no noticeable evidence of histologic injury by long-term CLEFMA treatment in mice.

**Figure 9:**
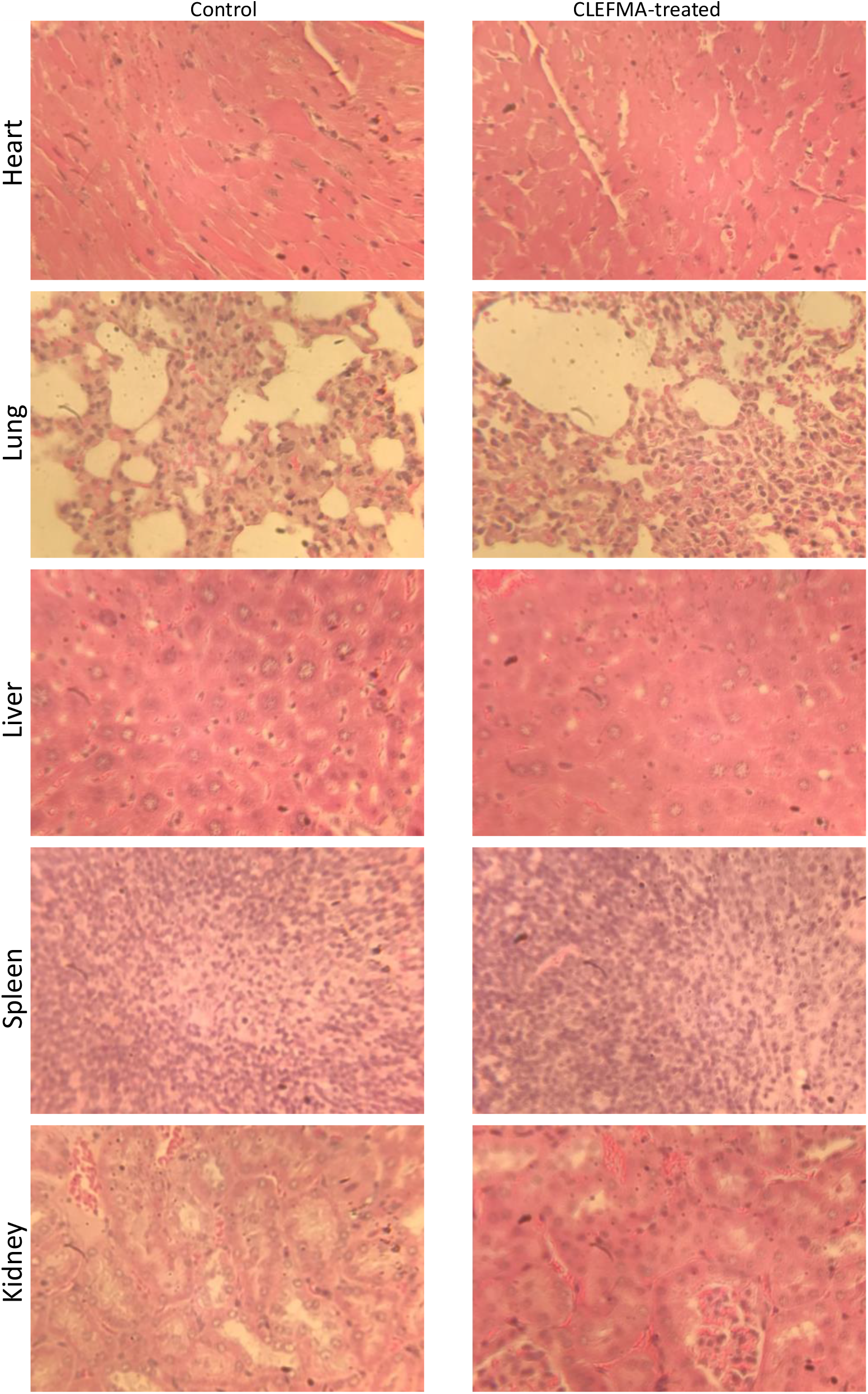
Effect of CLEFMA treatment on histology of heart, lung, liver, spleen, and kidney. The sections were H & E stained. The tissues were obtained from the mice recruited in 4-week/weekly-dosing study.

### Clinical chemistry, hematology, and urinalysis

The values of clinical chemistry parameters in mice treated with CLEFMA are shown in **Table 2**. From 1-week/daily-dosing study, only values for 50 mg/kg dose are given. Except for aspartate aminotransferase (AST), we did not find any significant difference between values from the untreated controls and those from CLEFMA-treated mice. However, since the AST levels were towards the lower side of control values, the possibility of liver damage by CLEFMA treatment could be rejected. The data included in **Table 3** also points out that, except for total WBCs, CLEFMA treatment did not significantly alter the hematological status as well. Similarly, urine samples showed no significant change after CLEFMA treatment (**Table 4**). The clinical chemistry, hematology, and urinalysis observations were similar in the 4-week/weekly-dosing study. These results suggest that CLEFMA treatment does not produce significant changes in clinical biochemistry parameters.

**Table 2.** Serum chemistry of healthy CD31 mice treated with CLEFMA.

| **Clinical Chemistry** | **1-Week/Daily-dosing** |  | **4-Week/Weekly-dosing** |  |
| --- | --- | --- | --- | --- |
| **Parameters** | **0 mg/kg** | **50 mg/kg** | **0 mg/kg** | **35 mg/kg** |
|  | **[n=4]** | **[n=3]** | **[n=4]** | **[n=4]** |
| **ALP [U/L]** | 79.0±26.9 | 84.0±23.5 | 49.0±17.0 | 23.0±8.0 |
| **ALT [U/L]** | 173.0±97.0 | 57.0±39.3 | 36.5±22.0 | 45.0±18.0 |
| **AST [U/L]** | 370.0±100.0 | 59.0±24.1* | 65.0±19.0 | 58.0±11.0 |
| **Albumin [gm/dL]** | 2.8±0.2 | 2.7±0.1 | 2.5±0.4 | 2.4±0.5 |
| **Globulin [gm/dL]** | 2.2±0.2 | 2.2±0.1 | 2.5±0.3 | 3.4±1.2 |
| **A/G Ratio** | 1.3±0.1 | 1.2±0.1 | 1.2±0.1 | 0.8±0.1 |
| **Total Protein [gm/dL]** | 4.9±0.3 | 4.8±0.1 | 5.4±0.5 | 6.2±0.8 |
| **Total Bilirubin [mg/dL]** | 0.1±0.05 | 0.1±0.05 | 0.1±0.05 | 0.1±0.05 |
| **BUN [mg/dL]** | 42.7±10.7 | 21.6±1.1 | 34.5±4.04 | 31.5±4.5 |
| **Creatinine [mg/dL]** | < 0.2 | < 0.2 | < 0.2 | < 0.2 |
| **Cholesterol [mg/dL]** | 119.0±45.0 | 106±17.6 | 86.5±39.0 | 103.0±7.2 |
| **Glucose [mg/dL]** | 200.0±46.2 | 238.0±31.2 | 179.0±77.0 | 296.0±54.0* |
| **Calcium [mg/dL]** | 9.6±0.4 | 9.6±0.4 | 10.3±0.3 | 10.5±0.5 |
| **Phosphorus [mg/dL]** | 7.4±2.3 | 7.5±0.5 | 8.4±1.4 | 6.5±0.5 |
| **Sodium [mmol/L]** | ND | ND | 159±5.5 | 157±2.3 |
| **Potassium [mmol/L]** | ND | ND | 5.8±1.0 | 5.0±0.1 |
| **Hemolysis** | No | No | No | No |

The values are given as mean ± SD (n = number of mice that survived and were tested for these parameters). ND = Not determined; AST=aspartate aminotransferase, ALT= alanine aminotransferase, ALP= Alkaline phosphatase, A/G=albumin to globulin ratio, BUN= blood urea nitrogen (\*p < 0.05 versus 0 mg/kg dose group)

**Table 3.** Hematology of healthy CD31 mice that received CLEFMA.

| **Hematology** | **1-Week/Daily-dosing** |  |
| --- | --- | --- |
| **Parameters** | **0 mg/kg** | **50 mg/kg** |
|  | **[n=1]** | **[n=3]** |
| **Total WBC (x10<sup>3</sup> cells/ μL)** | 14.0 | 7.1±0.4 |
| **RBC (10<sup>6</sup> cells/μL)** | 8.8 | 7.8±1.3 |
| **Hemoglobin (g/dL)** | 15.4 | 13.3±1.5 |
| **Hematocrit (%)** | 48.3 | 41.9±5.6 |
| **MCV (femtoliter/cell)** | 55.0 | 54.3±2.5 |
| **MCH (picogram/cell)** | 17.4 | 17.2±1.2 |
| **Neutrophils (x10<sup>3</sup> cells/ μL)** | 12.0 | 22.0±5.0 |
| **Lymphocytes (x10<sup>3</sup> cells/ μL)** | 83.0 | 70.0±6.9 |
| **Monocytes (x10<sup>3</sup> cells/ μL)** | 3.0 | 2.0±0.06 |
| **Eosinophils (x10<sup>3</sup> cells/ μL)** | 2.0 | 4.8±1.0 |
| **Basophils (x10<sup>3</sup> cells/ μL)** | 0 | 0.2±0.05 |
| **Polychromasia** | Slight | Slight |

The values are given mean ± SD (n = number of mice that survived and were tested for these parameters). WBC = white blood cell, RBC = red blood cells, MCV = mean corpuscular volume, MCH = mean corpuscular hemoglobin. In the dose group 0 mg/kg, sample from only one animal could be tested.

**Table 4.** Urine chemistry of healthy CD31 mice that received CLEFMA.

| **Urine** | **1-Week/Daily-dosing** |  | **4-Week/Weekly-dosing** |  |
| --- | --- | --- | --- | --- |
| **Parameters** | **0 mg/kg** | **50 mg/kg** | **0 mg/kg** | **35 mg/kg** |
|  | **[n=2]** | **[n=3]** | **[n=4]** | **[n=4]** |
| **Color** | Yellow | Yellow | Yellow | Yellow |
| **Clarity** | Clear | Clear | Clear | Clear |
| **Specific Gravity** | 1.08±0.06 | 1.1±0.01 | 1.04±0.0 | 1.05±0.0 |
| **pH** | 6.1±0.25 | 6.6±0.5 | 6.2±0.5 | 6.75±0.3 |
| **Glucose** | -ve | -ve | -ve | -ve |
| **Protein** | -ve | Traces | -ve | -ve |
| **Ketones** | Traces | Traces | -ve | -ve |
| **Urobilinogen** | Normal | Normal | Normal | Normal |
| **Bilirubin** | -ve | -ve | -ve | -ve |
| **Blood** | -ve | -ve | -ve | -ve |

The values are given as mean ± SD (n = number of mice that survived and were tested for these parameters).

## DISCUSSION

Treatment of advanced NSCLC patients largely relies on classical cytotoxic agents such as doxorubicin, cisplatin, and gemcitabine. Often cancer patients are either completely resistant to chemotherapy at the outset or gradually attain resistance over the course of therapy (Solyanik, 2010). To overcome drug resistance, a dose significantly larger than the usual dose is prescribed. However, most anticancer drugs are associated with inherent toxicity; the unintended exposure of normal cells to drugs leads to several undesired side effects. Even relatively benign drugs, such as celecoxib, possess cytotoxic effects in normal cells at the pharmacologic dose levels. Celecoxib is a selective COX-2 inhibitor which has been approved by the FDA for adjuvant therapy of colon and rectum polyps in patients with familial adenomatous polyposis [21, 22]. Unfortunately, at high concentrations needed for chemoprevention or cancer treatment, celecoxib also affects COX-2 independent targets, resulting in toxicity in normal tissues (Graham et al., 2005; Pyrko et al., 2008; Solomon et al., 2005). Our own data indicates that celecoxib affects the viability of normal lung fibroblasts at dose levels required to effectively kill lung cancer cells.

It is apparent that drugs with relative or absolute selectivity towards cancer cells are highly sought after in oncology. Here, we report the selectivity of anticancer activity of CLEFMA. CLEFMA is a chalcone-based molecule with a potent anti-proliferative activity [15]. The current understanding about the mechanistic basis of CLEFMA activity suggests that it perturbs redox homeostasis in cancer cells [16]. CLEFMA induces ROS production, sustained activation of extracellular signal-regulated kinase, and apoptotic cell death in lung adenocarcinoma H441 cells (unpublished data). Cancer cells are more sensitive to drug-induced generation of ROS because of their higher basal level of oxidative stress than the normal cells [23]. Recent research in anticancer drug development has amply demonstrated that chemicals that can induce moderate oxidative stress can trigger cancer cell apoptosis [24, 25] while sparing the normal cells from significant toxicity [26, 27]. Many existing anti-cancer drugs, that were initially thought to possess single cellular target, are now being found to work because of their ability to induce oxidative stress and generate ROS [28-31].

We investigated the effect of CLEFMA on cell viability and generation of ROS in multiple NSCLC and normal lung fibroblasts. CLEFMA induced cell death in all cancer cells in a dose-dependent manner, but did not affect the viability of normal cells. Further, CLEFMA-induced cell death was found to be mediated by induction of ROS which is generated selectively in cancer cells, but not in normal cells. Finally, CLEFMA was found to induce the expression of pro-apoptotic markers only in the cancer cells. Earlier we reported that CLEFMA-induced cell death is via caspase-mediated apoptotic pathway [18]. The pro-apoptotic effect of CLEFMA was typified by down-regulation of cell survival proteins such as cIAP1, Bcl2, Bcl-xL, and survivin as well as up-regulation of apoptosis inducers. Here, we chose to monitor caspase-3, PARP, and phospho-p53 as markers of apoptotic pathway and Bcl2, Bcl-xL, cIAP1, and survivin as anti-apoptotic proteins. CLEFMA treatment induced the cleavage of caspase 3 as well as PARP in cancer cells and increased the expression of phospho-p53 in cancer cells. At the same time, the expression levels of anti-apoptotic proteins were substantially reduced in CLEFMA-treated cancer cells. These alterations in expression of pro-apoptotic and anti-apoptotic markers were conspicuously missing in CLEFMA-treated normal lung fibroblasts.

Another typical feature of lung cancer development is its association with constitutively activated NF-κB pathway [32]. That CLEFMA inhibits NF-κB and the expression of its targeted genes has been previously reported [18]. Here we found that CLEFMA treatment in all three cancer cells inhibited the nuclear localization of phospho-p65 subunit of NF-κB. However, CLEFMA showed no effect on NF-κB phospho-p65 expression in the normal lung fibroblasts. Earlier, we reported that CLEFMA reduces the DNA-binding activity of NF-κB in H441 cancer cells, but not in normal fibroblasts CCL151 [18]. A variety of anticancer drugs inhibit the NF-κB pathway, either as their primary target or secondary to their effect on other pathways [33]. Non-specific and off-target effect of such drugs on NF-κB activity are potential causes of severe side effects, including debilitating immunosuppression. NF-κB plays a critical role in the maintenance of host defense responses and cell proliferation [34]. Considering the importance of NF-κB in maintaining normal growth and immunity, inhibition of NF-κB pathway for prolonged periods is not a recommended chemo preventive or therapeutic strategy. However, drugs like CLEFMA which provide selective NF-κB inhibition in cancer cells may able to overcome this issue. Although, our current understanding about the mechanistic basis of this observation is limited, we believe that the unique property of CLEFMA to inhibit NF-κB and induce redox stress is related to its proteasome-inhibiting activity. CLEFMA inhibits the 26S proteasome by interacting with regulatory subunit protein 13, an ubiquitin receptor [17, 35]. Proteasome inhibitors are known to suppress NF-κB signaling by interfering in degradation of inhibitor of κB [36]. The direct role of selective ROS production in CLEFMA-treated cells as the cause of NF-κB inhibition only in cancer cells also cannot be ruled out. ROS inhibit NF-κB [37-39], perhaps by oxidizing critical amino acid residues in the NF-κB subunits and thus interfering in its ability to bind DNA and influence transcription [40, 41]. Another explanation is based on the duration of ROS production. Accordingly, sustained ROS generation can inhibit NF-κB activity even though ROS can activate NF-κB initially [42].

The results discussed so far provided a strong basis of the usefulness of CLEFMA as a cancer cell-specific cytotoxic drug, although its low potency (IC_50_ approximately 5 µM) creates a doubt whether effective plasma concentration could be achieved. This raises a question of maximum dose of CLEFMA that could be administered, even when our previous studies showed that daily intraperitoneal dosing of 0.4 mg/kg bodyweight was effective in controlling tumor growth in rodent models [18, 43]. In a limited cohort of mice, we found that MTD_Survival_ and MTD_10%BW_ of CLEFMA were 10 and 50 mg/kg bodyweight, respectively. At an average dose of 35 mg/kg, no toxic effects of CLEFMA were observed in a 4-week/weekly-dosing study. In comparison, the reported MTD for cisplatin, gemcitabine, and doxorubicin in mice are 6 mg/kg, 120 mg/kg, and 3 mg/kg bodyweight, respectively [44-46]. Even though these drugs have more potent anticancer activity than CLEFMA, they do not possess selectivity towards cancer cells. The argument that ultra-potent drugs (IC_50_ in pico-to-nano Molar range) are better than the drugs with efficacy in low micro molar range is partially flawed, because potency without selectivity in action could often be counterproductive. Pharmaceutical wisdom dictates that if a drug is less potent but more selective, it should be the preferred one [47]. Secondly, anticancer drugs with one specific cellular target are candidates for the development of chemo-resistance. There is a wide agreement that current molecularly targeted approaches are not yielding the progress they promised. All cells have capacity to intelligently alter their biochemistry to survive when challenged with cytotoxic drugs and this capacity is visibly more in cancer cells. A drug that influences multiple pathways in cancer-specific manner could be an answer to the challenge of keeping cancer growth under control.

To the best of our knowledge, demonstration of this kind of selectivity at molecular levels of anti-cancer activity has not been a part of published literature. Although these results are limited to a few cancer and normal cell lines, the findings are encouraging. Secondly, since the MTD is 75 times more than the daily dose used for tumor suppression reported in our earlier study [18], a pre-clinical study using weekly dosing at 35 mg/kg is warranted.

## Acknowledgement

The work was supported by National Institute of Health R03 CA143614-01 and a Bridge research grant from Presbyterian Health Foundation, Oklahoma City (OK).

